# Bifunctional Peptide that Anneals to Damaged Collagen and Clusters TGF-β Receptors Enhances Wound Healing

**DOI:** 10.1101/2021.09.22.461420

**Authors:** Sayani Chattopadhyay, Leandro B. C. Teixeira, Laura L. Kiessling, Jonathan F. McAnulty, Ronald T. Raines

**Affiliations:** Department of Chemistry, School of Veterinary Medicine, University of Wisconsin–Madison, Madison, Wisconsin 53706, United States; Department of Pathobiological Sciences, School of Veterinary Medicine, University of Wisconsin–Madison, Madison, Wisconsin 53706, United States; Department of Biochemistry, School of Veterinary Medicine, University of Wisconsin–Madison, Madison, Wisconsin 53706, United States; Department of Surgical Sciences, School of Veterinary Medicine, University of Wisconsin–Madison, Madison, Wisconsin 53706, United States; Department of Chemistry, Massachusetts Institute of Technology, Cambridge, Massachusetts 02139, United States

## Abstract

Transforming growth factor-β (TGF-β) plays important roles in wound healing. The activity of TGF-β is initiated upon binding of the growth factor to extracellular domains of its receptors. We sought to facilitate activation by clustering these extracellular domains. To do so, we used a known peptide that binds to TGF-β receptors without diminishing their affinity for TGF-β. We conjugated this peptide to a collagen-mimetic peptide that can anneal to damaged collagen in a wound bed. We find that the conjugate enhances collagen deposition and wound closure in mice in a manner consistent with the clustering of TGF-β receptors. This strategy provides a means to upregulate the TGF-β signaling pathway without adding exogenous TGF-β and could inspire means to treat severe wounds.

**TOC Graphic:** 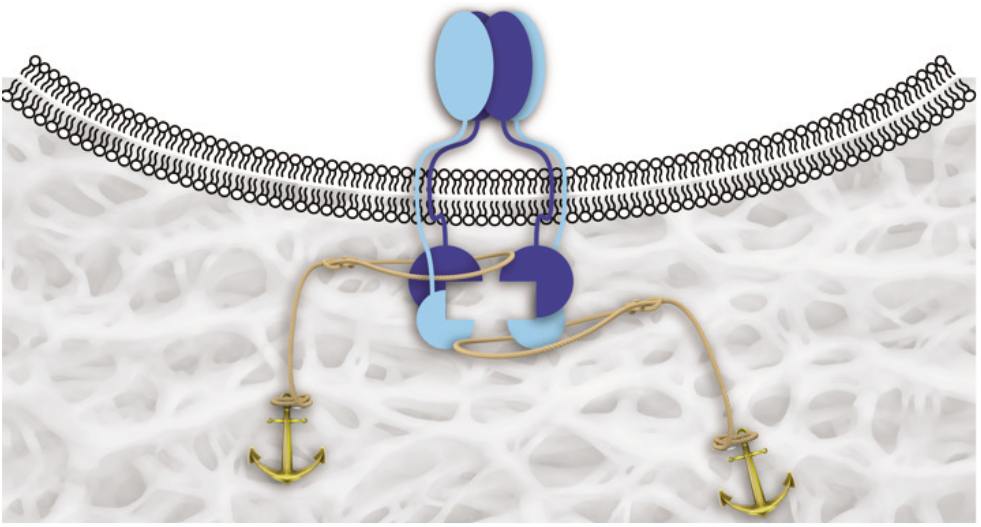

## INTRODUCTION

The transforming growth factor β (TGF-β) family is a group of mammalian secretory proteins that play myriad roles in development and disease.^1–4^ Their biological activities are initiated upon interaction with two type-I receptors (TβRI) and two type-II receptors (TβRII).^5–7^ These cell-surface receptors are characterized by an extracellular TGF-β–binding domain, a transmembrane domain, and a cytosolic serine/threonine kinase domain.^8^ Upon binding to TGF-β, TβRII recruits TβRI into an activated heterotetrameric complex (Figure 1A). The cytosolic domain of TβRII then catalyzes the phosphorylation of the regulatory region of TβRI and thereby activates the adjacent serine/threonine kinase domain, leading ultimately to the phosphorylation of downstream effectors, Smad2 and Smad3, which translocate to the nucleus and mediate gene expression.

**Figure 1.**
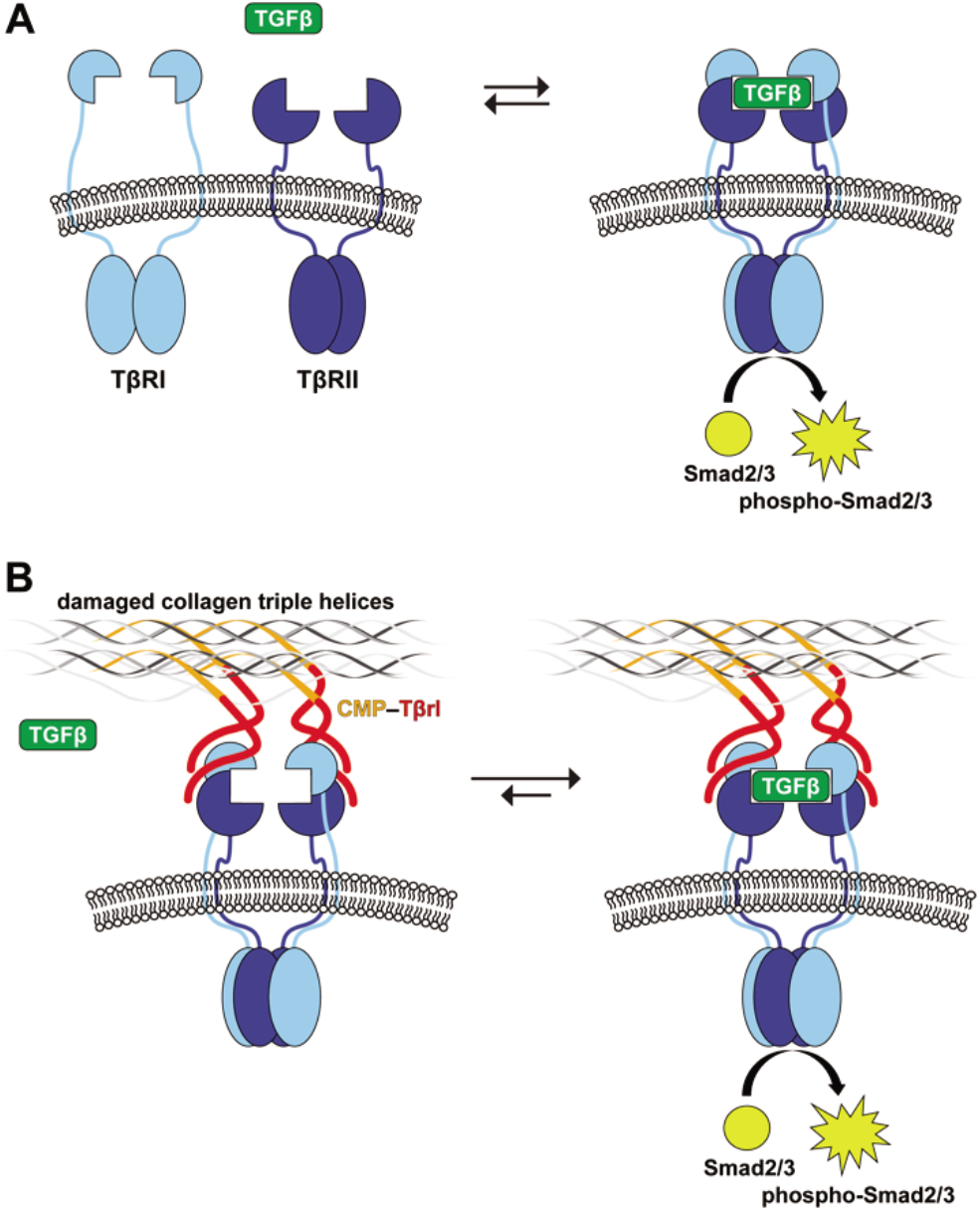
Representation of TGF-β receptor–ligand complex formation and its activation of the Smad2/3 proteins by their cytosolic kinase domains. (A) TGF-β induces the heterotetramerization of type-I and type-II receptors (TβRI and TβRII, respectively). (B) Clustering of TβRI and TβRII by binding to a ligand (**Tβrl**) that is immobilized in a wound bed upon annealing of a pendant **CMP**.

Activation of the TGF-β signaling cascade can promote dermal fibrosis.^9–13^ For example, TGF-β administered to full-thickness wounds in rabbits by encapsulation in a collagen-sponge scaffold accelerates re-epithelialization and contraction.^14^ Similarly, TGF-β applied topically or by injection heals wounds by increasing tensile strength and promoting fibroblast proliferation and collagen deposition.^15–20^ These approaches, however, require the administration of exogenous TGF-β, which can have deleterious consequences.^9–12^

Using phage display, we identified dodecapeptides that bind to the extracellular domains of both TβRI and TβRII to form complexes with *K_d_* ≈ 10^−5^ M.^21^ These peptides do not, however, antagonize the binding of TGF-β. In addition, we demonstrated that a multivalent display of these ligands on a PEG-based dendrimer increases their functional efficacy for the receptors.^21^ Likewise, immobilization of the ligands on a synthetic surface enables activation by subpicomolar concentrations of endogenous TGF-β.^22^ Gene expression profiles revealed that the surfaces regulate TGF-β responsive genes selectively. Now, we sought to exploit these ligands in vivo.

Collagen is the major component of the extracellular matrix.^23–25^ We and others have reported on the use of collagen-mimetic peptides (CMPs) to anchor probes and growth factors in damaged or abnormal collagen.^26–34^ Here, we use a CMP conjugate to immobilize a peptidic ligand for TGF-β receptors in wound beds of mice. Our intent is to cluster cell-surface TβRI and TβRII and thereby enhance the sensitivity of cells to circulating TGF-β (Figure 1B). As TGF-β is involved in various stages of wound healing,^9–12^ we validate the efficacy of our approach by using a variety of assays.

## RESULTS AND DISCUSSION

### Peptide Design

As an effector to sensitize cell-surface receptors to TGF-β signaling, we chose to use the LTGKNFPMFHRN peptide.^21^ As a CMP, we chose (PPG)_7_ because of its simplicity and demonstrated efficacy in relevant contexts in vitro, ex vivo, and in vivo.^26,28,29,32,33^ In its selection, LTGKNFPMFHRN (**Tβrl**) was displayed as a fusion to the N-terminus of the PIII coat protein of phage.^21^ We mimicked this display by conjugating **Tβrl** to the N-terminus of **CMP**. Accordingly, we synthesized the 33-mer peptide **Tβrl–CMP** and its **Tβrl** and **CMP** components by solid-phase peptide synthesis (SPPS).

### Mouse Model

Wound healing is a complex process. Mouse models have illuminated the mechanisms that underlie wound healing and established translatable therapeutic strategies.^35–38^ We chose to use diabetic (*db/db*) mice as our model. These mice exhibit characteristics similar to adult human onset type II diabetes mellitus,^39,40^ including impaired wound healing.^41^ Excisional wounds in *db/db* mice show a delay in wound closure, decreased granulation tissue formation, decreased vascularization in the wound bed, and diminished cell proliferation.^42^ The course of wound healing in these mice follows closely the clinical observations of human diabetic patients.^43^ For example, these mice show delayed and reduced expression of keratinocyte growth factor and peripheral neuropathy,^44^ as do diabetic humans.^45^

We chose to use an excisional wound model, which heals from the wound margins and provides the broadest assessment of the various parameters for wound healing, such as re-epithelialization, fibrovascular proliferation, contracture, and angiogenesis.^36^ This wound model also provides for large dorsal surfaces that simplify application of topical agents directly into the wound bed, as well as the availability of two wounds side-by-side on the same mouse.

### Unsplinted Wounds

Wounds were created in the craniodorsal region of *db*/*db* mice under anesthesia and treated topically with **Tβrl-CMP** (25 μL of a 20 mM solution in 5% PEG/saline solution).^46,47^ **CMP** and **Tβrl** (25 μL of 20 mM solutions in 5% PEG/saline) were also tested individually. The delivery vehicle itself was tested as a control.

Fibrovascular influx and the deposition of new collagen in wounds were measured by examining the picosirius red-stained histologic sections under polarized light^48^ and expressed as a percentage of the total wound area. The picosirius red stain highlights areas of new collagen deposition, as well as extant dermal collagen. Due to more extensive crosslinking and maturation, older collagen is stained more brightly and densely in comparison to newly formed collagen. Upon its release from degranulating platelets, TGF-β1 can attract fibroblasts chemotactically to a wound site^49–51^ and stimulate their proliferation.^52^ As part of a positive feedback mechanism, fibroblasts release additional TGF-β in response and promote collagen biosynthesis.

Compared to control wounds treated with **Tβrl**, **CMP**, or the delivery vehicle, wounds exposed to **Tβrl–CMP** exhibited a significant increase in the amount of collagen deposited in the wound bed. (Figure 2A). This result is consistent with reports in which the topical application of TGF-β in animal models enhanced the production of collagen and fibronectin by fibroblasts^53,54^ and potently stimulated granulation tissue formation in wound-healing models.^55,56^ In contrast, the production of collagen was diminished in the presence of anti–TGF-β antibodies.^57^

**Figure 2.**
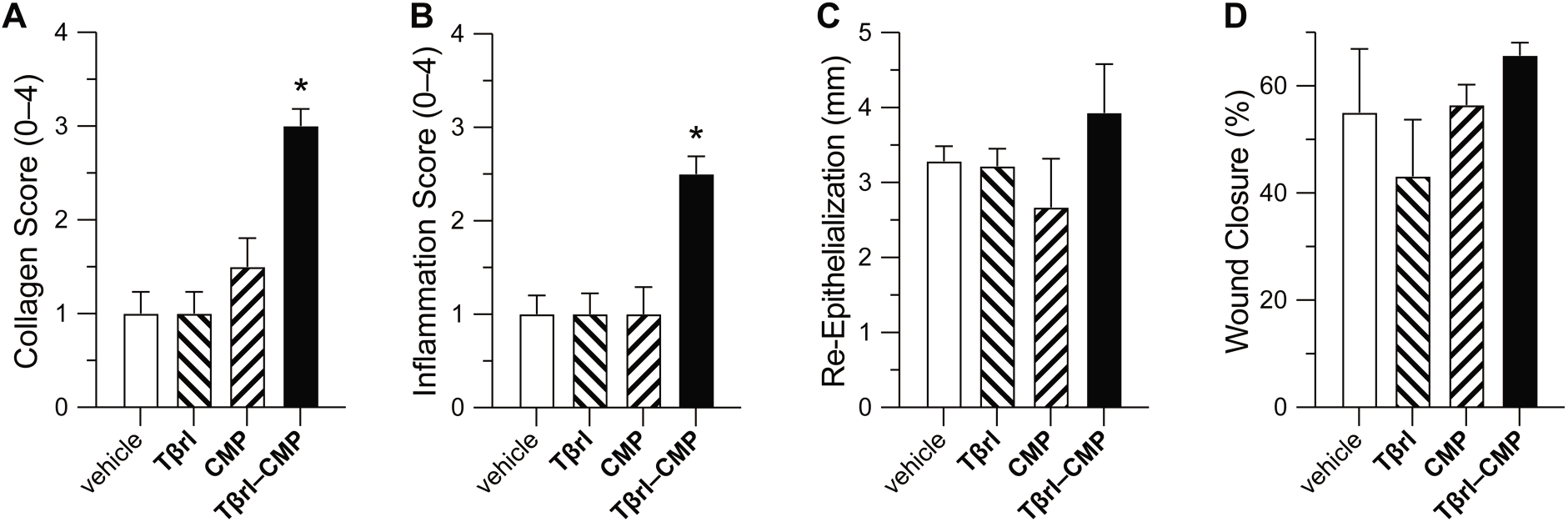
Effect of **Tβrl–CMP** (0.5 μmol) and controls on the healing of unsplinted cutaneous wounds in mice. Values are the mean ± SE (*n* = 10 wounds in 5 mice) with \**p* < 0.05. (A) Fibrovascular influx in wounds on a scale of 0–4 by Day 12 post-surgery. (B) Inflammation in wounds on a scale of 0–4 on Day 12 post-surgery. (C) Re-epithelialization of wounds by Day 16 post-surgery. (D) Wound closure by Day 12 post-surgery, calculated as the wound size as a percentage of the original wound size on Day 0.

The wound-healing process is associated with a transient accumulation of fibroblasts that express elevated levels of TβRI and TβRII.^15–20^ The highest cellular density is observed in the deepest regions of the granulation tissue.^58^ **Tβrl–CMP** tethered to the wound bed is poised to preorganize these receptors and thereby enhance the cellular sensitivity to TGF-β signaling and consequent formation of new collagen, without a need for exogenous TGF-β (Figure 1). Earlier work showed that tethering TGF-β to a PEG-based polymer scaffold caused a significant increase in matrix production and collagen deposition.^59^ Such a treatment also counteracted the attenuation of ECM production, which is observed otherwise in the presence of biomaterials containing cell-adhesive ligands.^60^ The **Tβrl-CMP** conjugate can behave in a similar manner and promote cellular adhesion while strengthening the wound bed itself by enhancing collagen synthesis and consequent ECM production. **Tβrl** alone cannot, however, access such preorganization and provides a response that is indistinguishable from the delivery vehicle.

A similar outcome was apparent when wounds were analyzed for an inflammatory response. The concentration of TGF-β is ~1 pM in human serum.^61^ Upon cutaneous injury, TGF-β levels elevate rapidly.^62,63^ For example, TGF-β levels reach a peak at 3 days post 6-mm full-thickness wounding in transgenic mice, which coincides with the peak of the inflammation during early stages of wound healing.^63^ Subcutaneous injection of TGF-β affords a histological pattern for neutrophil and macrophage recruitment, fibroblast proliferation, and vascular growth, similar to the process of normal inflammation and repair in cutaneous wounds.^55^ In early stages, TGF-β provides a highly chemotactic ligand for human peripheral blood monocytes,^64^ which is critical for the initiation of an inflammatory response. By a positive feedback mechanism, the recruited monocytes and macrophages produce more TGF-β (thereby perpetuating their activity) as well as mitogenic and chemotactic substances that act on other cells. Wounds treated with **Tβrl-CMP** showed an inflammatory influx that is significantly greater than that of **Tβrl**, **CMP**, or the delivery vehicle (Figure 2B). The macrophages once activated or during maturation, downregulate their receptors for TGF-β and hence their ability to be stimulated any further.^64^ The peripheral blood monocytes also become susceptible to deactivation by TGF-β,^65^ which almost self-regulates the inflammatory stage by inhibiting the proteolytic environment created by the inflammatory cells and easing the healing process into the proliferative phase.^66^

Re-epithelialization is the process by which keratinocytes both proliferate and migrate from wound edges to create a barrier over the wound. The role of TGF-β in this process is not completely understood. In vitro, TGF-β inhibits the proliferation of keratinocyte but enhances their migration.^67,68^

Data in vivo are contradictory. Transgenic mice that overproduced TGF-β show enhanced epithelialization in partial-thickness wounds^69^ and anti–TGF-β antibodies administered to rabbits impairs epithelialization.^70^ On the other hand, mice null for Smad3 show accelerated keratinocyte proliferation and epithelialization upon administration of TGF-β compared to wild-type mice.^50,71^ In our experiment, we observed a tendency towards increased epithelialization of wound beds treated with **Tβrl-CMP** (Figure 2C). This result is consistent with cells responding quickly to endogenous TGF-β, which modulates the proliferative and migratory properties of the keratinocytes.^67,68^

No discernible differences were apparent in the rate of wound closure in the treated wounds (Figure 2D). This equivalence could be due to the preponderance of wound closure by contracture in rodents. That obscures wound changes related to epithelial closure or from the formation of scabs, removal of which disturbs the newly formed epidermis and could make wound size measurements inconclusive. We therefore took an alternative, complementary experimental approach.

### Splinted Wounds

Mouse models are informative but dissimilar from human skin models in that the major mechanism of wound closure is contraction; in humans, re-epithelialization and granulation tissue formation are the major phases.^35–38^ The use of splints around excisional wounds in mice forces healing to occur by granulocyte formation and re-epithilialization, while minimizing the effects of contraction compared to un-splinted wound models.^72^ As described previously,^28^ we sutured silicone O-rings around the wound margins to act as splints. We increased the cohort size to eight mice per group. Wounds were treated with 25 μL of **Tβrl-CMP**, **Tβrl**, **CMP**, or the delivery vehicle, as with the unsplinted wounds. The wounded mice were then allowed to recover and monitored over a period of 16 days.

Histopathological analysis post-euthanasia on Day 16 revealed that 15 of the 16 wounds treated with **Tβrl-CMP** were closed completely (Figure 3). Likewise, the wound size was significantly lower than that for the delivery vehicle (Figure 4A). Treatment with **Tβrl** also showed slightly enhanced wound closure, though it was not significantly different from the delivery vehicle and comparable to treatment with **CMP**. Upon comparing the extent of re-epithelialization, we noticed a clear tendency of improved keratinocyte proliferation in the wound beds treated with **Tβrl-CMP** in comparison to wounds treated with **CMP** or the delivery vehicle (Figure 4B). TGF-β has been reported to promote epithelial cell attachment and migration in vivo^73,74^ and to stimulate the expression of keratinocyte integrins during re-epithelialization.^75^ Keratinocyte migration takes place across a substrate, typically the dermis. The deposition of a substantial granulation tissue layer in the longer time period of the splinted-wound experiments (16 days), could have provided the requisite surface for the migration of keratinocytes and increased the length of newly formed epithelial layer.

**Figure 3.**
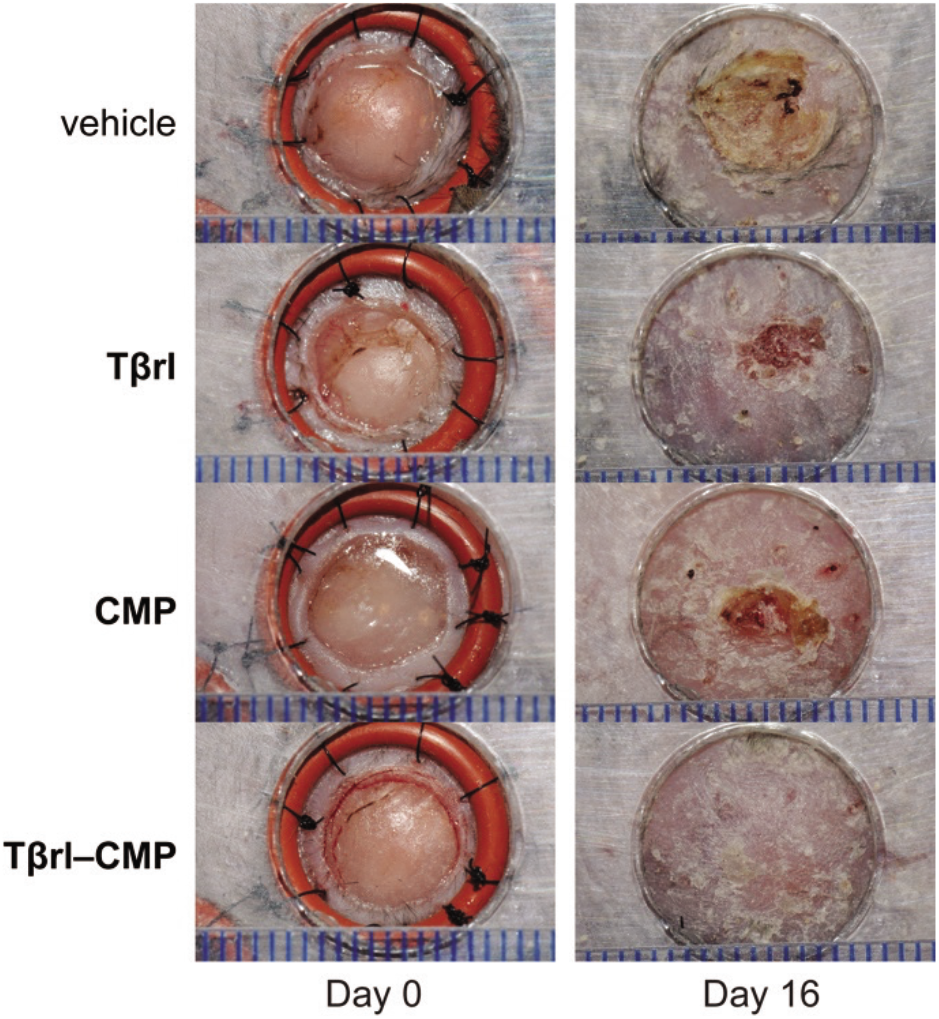
Representative images of splinted wounds in mice on Day 0 (*i.e*., immediately post-surgery) and on Day 16 post-surgery after removal of splints but before euthanasia. Wounds were treated with vehicle (5% PEG/saline), **Tβrl** (0.5 μmol), **CMP** (0.5 μmol), or **Tβrl-CMP** (0.5 μmol). In mice treated with **Tβrl-CMP,** 15 of 16 wounds showed complete closure. Scale bars (blue) are separated by 1.0 mm.

**Figure 4.**
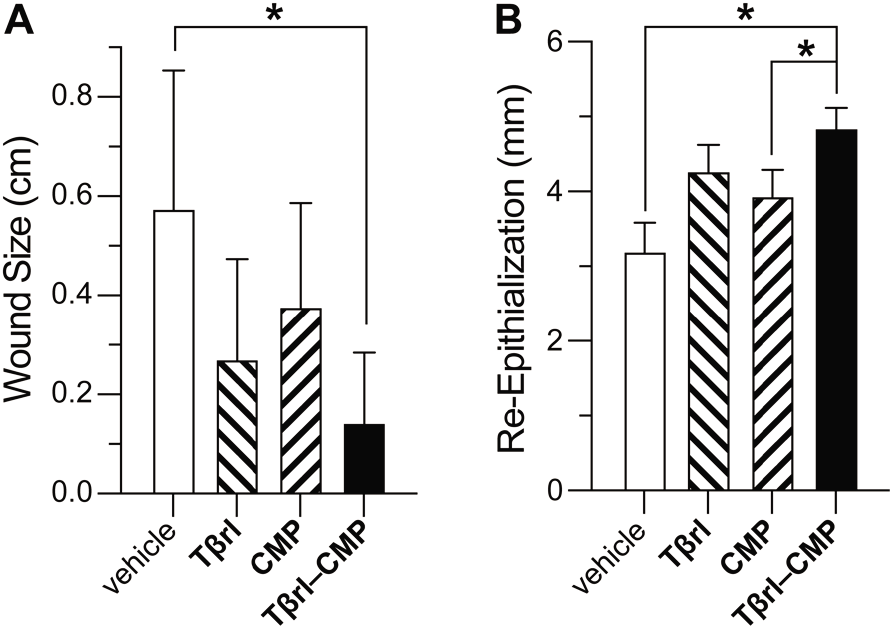
Effect of **Tβrl-CMP** (0.5 μmol) and controls on the healing of splinted cutaneous wounds in mice. (A) Wound closure on Day 16 post-surgery, represented as the distance between advancing edges at the widest diameter. (B) Re-epithelialization of wounds by Day 16 post-surgery. Values are the mean ± SE (*n* = 16 wounds in 8 mice) with \**p* < 0.05.

### Dose–Response Analyses

Finally, we assessed the potency of the TGF-β receptor ligand in ECM formation by treating the wounds with increasing doses of **Tβrl–CMP**. Identical 6-mm o.d. wounds were created on the backs of *db/db* mice (5 mice/10 wounds per group), and then treated with a 25-μL solution of **Tβrl-CMP** (0.080–50 mM) for 30 min. The mice were allowed to recover, and the wounds were analyzed after a period of 12 days. The amount of newly formed collagen in the wound bed was identified with picosirius red at a depth of 0.75 mm from the healed surface.

The extent to which collagen was deposited were comparable in the stain and expressed as a percentage of marked area in the wound wounds treated with 0.08, 0.4, 2.0, and 10.0 mM solutions but increased in wounds treated with 50 mM **Tβrl-CMP** (Figure 5A). These data were mirrored in histopathological analyses (Figure 6).

**Figure 5.**
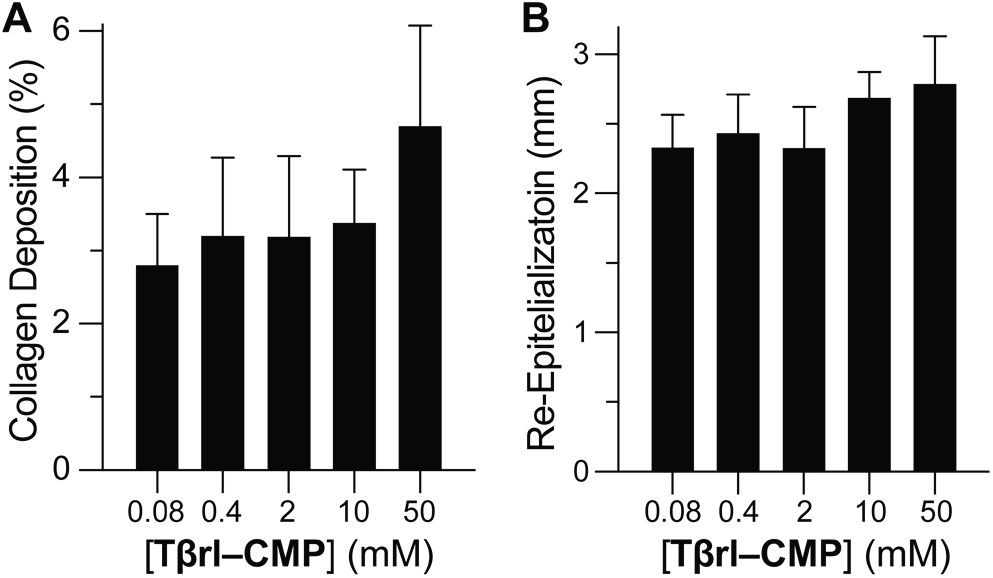
Dose–response analysis of unsplinted cutaneous wounds in mice to **Tβrl-CMP** and controls. Wounds were treated with a 25-μL solution of **Tβrl-CMP** in 5% PEG/saline and analyzed on Day 12 post-surgery. Values are the mean ± SE (*n* = 16 wounds in 8 mice). (A) New collagen deposition at a depth of 0.75 mm from the healed surface. (B) Re-epithelialization of wounds.

**Figure 6.**
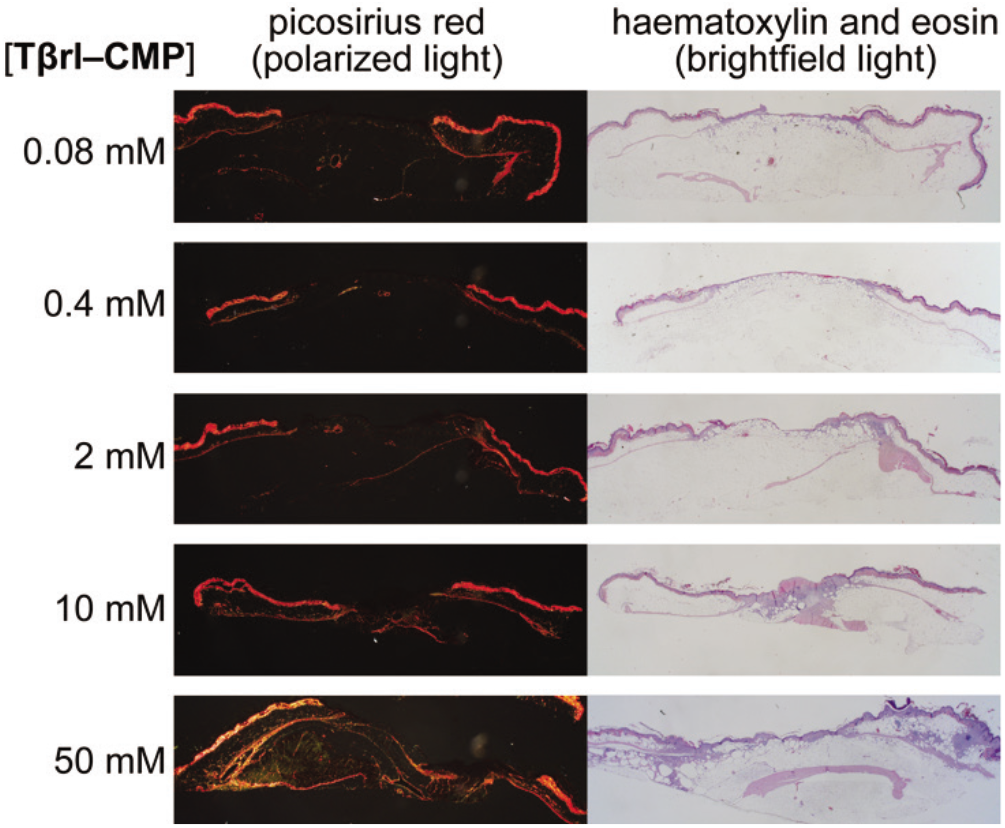
Representative histological images of **Tβrl–CMP**-treated wounds on Day 12 post-surgery. Wounds were stained with picosirius red and imaged with polarized light or with hemotoxylin and eosin and imaged with brightfield light.

Wound re-epithelialization did not exhibit a marked dependence on **Tβrl-CMP** dose, though there seemed to be a tendency for a higher response upon treatment with higher doses (Figure 5B). The inflammatory response showed the presence of mononuclear cells in larger amounts in wounds treated with higher doses in comparison to lower doses, which retained discernible amounts of neutrophils and polymorphonuclear cells.

## CONCLUSIONS

A **Tβrl-CMP** conjugate can upregulate collagen formation in mice. This result along with in vitro data^21,22^ are consistent with its clustering of TGF-β receptors and thereby sensitizing cells to endogenous TGF-β. This mode of action is unique for a pendant on a collagen-mimetic peptide that is annealed to a wound bed and provides opportunities. The closure of severe wounds where viable tissue has been destroyed by trauma involves the deposition of new collagen in accord with the severity of the damage. Amplified TGF-β activity during initial stages of the wound healing process stimulates fibroblast proliferation and activity, as well as keratinocyte migration over the surface of the wounds. This action accelerates the closure of wounds and the acquisition of tensile strength. Dose–response studies suggest that the amount of collagen deposition can be regulated without substantial changes in the rate of epithelialization. Although this type of healing could elicit scar formation but could provide an effective means to treat severe damage, as incurred from 3^rd^- or 4^th^-degree burns or traumatic mechanical damage, which might otherwise lead to lifelong impairment.

## EXPERIMENTAL PROCEDURES

### Materials

Commercial chemicals were of reagent grade or better, and were used without further purification. Anhydrous solvents were obtained from CYCLE-TAINER^®^ solvent delivery systems from J. T. Baker (Phillipsburg, NJ). HPLC grade solvents were obtained in sealed bottles from Fisher Chemical (Fairlawn, NJ). In all reactions involving anhydrous solvents, glassware was either oven- or flame-dried. Polyethylene glycol 8000 (PEG) from Fisher Chemical and bacteriostatic saline (0.9% w/v sodium chloride) from Hospira (Lake Forest, IL) were used to prepare 5% w/v PEG in saline solution as a delivery vehicle for treatments.

Male mice (*n* = 92) (BKS.Cg-*Dock7^m^* +/+ *Lepr^db^*/J) were from Jackson Laboratory (Bar Harbor, ME). Isoflurane was from Abbott Laboratories (Abbott Park, IL), buprenorphine·HCl was from Reckitt Benckiser (Berkshire, UK), and chlorhexidine gluconate (4% w/v) was from Purdue Products (Stanford, CT). Silicone O-rings (15 mm o.d., 11 mm i.d., 2 mm thickness) were from McMaster Carr (Chicago, IL).

### Instrumentation

Semi-preparative HPLC was performed with a Varian Dynamax C-18 reversed-phase column. Analytical HPLC was performed using a Vydac C-18 reversed-phase column. Mass spectrometry was performed with an Applied Biosystems Voyager DE-Pro matrix-assisted laser desorption/ionization mass spectrometer from Life Technologies (Carlsbad, CA) in the Biophysics Instrumentation Facility at University of Wisconsin-Madison.

### Peptide Synthesis and Purification

Peptides were SPPS using a 12-channel Symphony peptide synthesizer from Protein Technologies (Tucson, AZ) at the University of Wisconsin-Madison Biotechnology Center. Peptides were synthesized on Novabiochem FmocGly-Wang resin (0.4-0.7 mmol/g, 100–200 mesh) from EMD Chemicals (Gibbstown, NJ). Amino acids were converted to active esters by treatment with 1-hydroxybenzotriazole (HOBt, 3 equiv), *O*-benzotriazole-*N,N,N’,N’*-tetramethyl-uronium-hexafluoro-phosphate (HBTU, 3 equiv), and *N*-methylmorpholine (NMM, 6 equiv). Fmoc-deprotection was achieved by treatment with piperidine (20% v/v) in DMF.

**CMP** and **Tβrl-CMP** were synthesized by the sequential coupling of FmocProOH and FmocProProGlyOH. The first proline residue in was coupled to the resin after a swell cycle, and the next 7 residues were installed by using excess (5 equiv) FmocProOH, FmocProProGlyOH (which was synthesized as reported previously^76^), FmocAsn(Trt)OH, FmocArg(Pbf)OH, FmocHis(Trt)OH, FmocPheOH, FmocMetOH, FmocLys(Boc)OH, FmocGlyOH, FmocThr(*t*Bu)OH, and FmocLeuOH. **CMP** was cleaved from the resin by using 95:2.5:2.5 trifluoroacetic acid/triisopropylsilane/water (total volume: 2 mL); **Tβrl-CMP** was cleaved from the resin by using 92.5:5:2.5 trifluoroacetic acid /thioanisole/ethanedithiol (total volume: 2 mL). Both peptides were precipitated from *t*-butylmethylether at 0 °C, isolated by centrifugation, and purified by semi-preparative HPLC using linear gradients: **CMP**, 5–85% v/v B over 45 min; **Tβrl-CMP**, 10–90% v/v B over 50 min. Solvent A was H_2_O containing TFA (0.1% v/v); solvent B was CH_3_CN containing TFA (0.1% v/v). **CMP** was readily soluble in water but **Tβrl-CMP** required the addition of CH3CN (20% v/v) to form a clear solution for HPLC analysis. All peptides were judged to be >90% pure by HPLC and MALDI–TOF mass spectrometry: (*m*/*z*) [M + H]^+^ calcd for **CMP**, 1777; found 1777; (*m*/*z*) [M + H]^+^ calcd for **Tβrl-CMP**, 3221; found 3221.

**Tβrl** was either obtained from Biomatik (Wilmington, DE) or synthesized at the Peptide Synthesis Facility on Fmoc-Asn(Trt)-Wang resin (0.54 mmol/g, 100–200 mesh, Novabiochem^®^, EMD Chemicals, Gibbstown, NJ).

### Mouse Models

Male mice (homozygous for *Lepr^db^*, Jackson Laboratories, Bar Harbor, ME) aged 8-12 weeks were housed in groups until the day of surgery and then in separate cages post-surgery. The experimental protocol followed was according to the guidelines issued by the Institutional Animal Care and Use Committee (IACUC) at the University of Wisconsin-Madison. Mice were provided food and water *ad libitum*, as well as enrichment, and housed in a temperature-controlled environment with 12-h light and dark cycles.

On the day of the surgery, mice were anaesthetized with isoflurane gas using an induction chamber. For pain management, buprenorphine·HCl (0.01 mg/mL in 0.9% w/v saline) was injected subcutaneously (0.4 mL per mouse). Eyes were lubricated, and hind nails were clipped. The craniodorsal region was shaved using electric clippers, and the shaved area was scrubbed with alternating cotton swabs of chlorhexidine and sterile saline in circular strokes. Residual hair was removed. For the non-splinted wound model, identical 8-mm wounds were created on each side of the body with a biopsy punch, and the wounding was completed using forceps and scissors to prevent the punch from lacerating the subcutaneous tissue. The wounds were treated with **Tβrl-CMP** or controls, and then allowed to incubate for 30 min while the mouse was still under anesthesia. Wounds were photographed and the mice were then allowed to recover in their cages. For the dose–response experiments, un-splinted wounds were created with a 6-mm biopsy punch and then treated with 25 μL of **Tβrl-CMP** in 5fold increase of concentration, followed by incubation for 30 min under anesthesia.

For the splinted-wound model, splints were bilaterally placed in a symmetrical arrangement using adhesive and then secured to the skin using 8 interrupted 5-0 nylon sutures, encircling the splints with the knots.^77^ Wounds were created in the center of the splints using an 8-mm biopsy punch, and then removing the skin using forceps and scissors. The wounds were then treated with **Tβrl-CMP** or controls, and allowed to incubate for 30 min while the mouse was still under anesthesia. The wounds were photographed, and the mice were allowed to recover on a warming pad. ImageJ software^78^ from the National Institutes of Health (Bethesda, MD) was used to calculate the wound area (mm^2^) from digital photographs. Wound closure was defined as the reduction in area between wound edges over the course of the study and was reported as a percentage of the original wound size.

Mice were monitored daily for behavioral changes, and their body weights were recorded on days 1, 3, 6, 9, 12, and 16. The splints were checked daily, and any broken or untied suture was replaced according to the experimental protocol (*vide supra*). If only one suture was compromised during a 24-h period, it was replaced with a new suture. If, however, two or more sutures were compromised during a 24-h period, the wound was no longer considered splinted and was removed from the study.

### Wound Harvesting

Histopathology cassettes were labeled for mouse and wound identification. Note cards (1 inch × 1 inch square) were fitted to the bottom of the histopathology cassettes, and one edge was labeled “cranial” and aligned with the cranial side of the harvested wound. On the final day of the experiment, mice were euthanized with Beuthanasia^®^-D (0.5 mL per mouse). Using a scalpel blade and scissors, a ¾ inch × ¾ inch square area of tissue was taken from the mouse, keeping the wound centered in the tissue section. Deep dissection was performed to harvest several layers of tissue deep in the wound. The square section of tissue was affixed to the note card, with the cranial edge aligned with the labeled edge of the card. The cassettes were then closed and placed in formalin-filled jars for histopathological analysis.

### Histopathological Analyses

After euthanasia, the entire wound bed as well as the intact skin margin >5 mm was excised to the retro-peritoneum. The harvested tissue was then fixed in 10% v/v formalin for at least 24 h, and then sectioned through the center of the lesion. The center was marked with India ink prior to fixation. Routine paraffin processing was performed, and the tissue samples were sectioned serially at a thickness of 5 μm, ensuring that the center of the lesion was included on the slide. The slides were then stained with hematoxylin, eosin, and picosirius red. Sections were photographed under a light microscope with a mounted DP72 digital camera from Olympus (Shinjuku, Tokyo, Japan). The size of the wound, length of re-epithilialization, amount of fibrovascular proliferation in the dermis, and inflammatory response were measured on the slides containing the center of the lesion, and images were analyzed with CellScience Dimension 1.4 software from Olympus. The size of the wound was defined as the area of the wound not covered by an advancing epithelial layer and was calculated by measuring the distance between the opposite free edges of the wound. The length of re-epithelialization was defined as the length of the layer of proliferating keratinocytes covering the wound area and was calculated by measuring the distance between the free edge of the keratinocyte layer and the base where the cells were still associated with native dermal tissue. Both sides of the lesion were measured, and the final result was the sum of the two measurements. For wounds that had undergone complete re-epithelialization, a single measurement was taken from base to base.

Fibrovascular dermal proliferation was measured by examining the picosirius red-stained sections under polarized light, which highlighted the newly deposited dermal collagen. CellScience Dimension 1.4 software was used to select the wound bed; the amount of new collagen in the selected area was measured and expressed as a percentage of the total wound area or with a semi-quantitative histopathological 0–4 scoring system, where 0 indicated no discernible collagen formation, 1 indicated that <25% of the wound area was covered with fresh collagen, 2 indicated that 25–50% of the wound area was covered with fresh collagen; 3 indicated that 50–75% of the wound area was covered with fresh collagen, and 4 indicated that >75% of the wound area was covered with fresh collagen. The inflammatory response was assessed with a semi-quantitative histopathological 0–4 scoring system, where 0 indicated no inflammation, 1 indicated that <25% of the wound area was affected, 2 indicated that 25–50% of the wound area was affected, 3 indicated that 50–75% of the wound area was affected, and 4 indicated that >75% of the wound area was affected. The inflammatory response was also categorized as “acute” when >75% of the cells were neutrophils; “chronically active” when there was a 1:1 ratio of neutrophils and mononuclear cells; and “chronic” when >75% of the inflammatory cells were mononuclear.

### Statistical Analyses

All data were analyzed with a Mann–Whitney rank sum test, and statistical significance was set to *p* < 0.05. Statistical analyses were executed with Prism Version 5.0 software from GraphPad Software (La Jolla, CA).

## Funding

This work was supported by grants RC2 AR058971, R56 AR044276, R01 AI055258, and R01 GM049975 (NIH). MALDI–TOF mass spectrometry was performed at the University of Wisconsin-Madison Biophysics Instrumentation Facility, which was established with Grants BIR-9512577 (NSF) and S10 RR013790 (NIH).

## Notes

The authors declare no competing financial interest.

## ACKNOWLEDGMENTS

We are grateful to Lingyin Li and Nicholas L. Abbott for their guidance and advice. We thank Patricia Kierski, Diego Calderon, Dana Tackes, Kevin Johnson, and Zachary Joseph for help with the wound surgery and animal care.

## ABBREVIATIONS

CMP: collagen-mimetic peptide (here, (PPG)_7_)
TGF-β: transforming growth factor-β
CMP–Tβrl: collagen-mimetic peptide-transforming growth factor-β receptor ligand conjugate (here, LTGKNFPMFHRN-(PPG)_7_)
TβRI: transforming growth factor-β type-1 receptor
TβRII: transforming growth factor-β type-2 receptor
Tβrl: transforming growth factor-β receptor ligand (here, LTGKNFPMFHRN)

## Notes

### Competing Interest Statement

The authors have declared no competing interest.

